# Learned associations constrain anticipatory visual dynamics and enhance sensory and motor readiness to facilitate perceptual decisions

**DOI:** 10.64898/2026.11.10.765786

**Authors:** Lei Zhang, Mohan Yuan, Haoyu Zhang, Morris Moscovitch, Claude Alain

## Abstract

Learned cue–target associations can guide attention to facilitate perceptual decision-making, yet how they shape the content and dynamics of neural activity, and whether this facilitation reflects anticipatory preparation, post-target sensory enhancement, or both, remains unclear. Here, we used magnetoencephalography while participants made living/non-living judgments about visual objects (e.g., cat/violin) with left-or right-hand button presses. Cue–category associations were established through repeated pairings: predictive cues always preceded one target category, whereas uninformative cues preceded both categories equally often. Predictive cues facilitated behavioral responses despite limited explicit awareness of the learned associations. Representational similarity analysis showed that prediction preactivated target-specific representations in the visual cortex resembling target-evoked activity before target onset. Prediction also elicited anticipatory, response-specific motor beta suppression. Together, these findings supported anticipatory preparation, with little evidence of enhanced post-target visual specificity. Post-target visual specificity and anticipatory motor preparation independently predicted response speed, with hierarchical drift-diffusion modeling linking greater visual specificity to faster evidence accumulation and stronger motor preparation to shorter non-decision time (a model parameter reflecting sensory encoding and motor execution). Beyond representational content, prediction also constrained the evolution of anticipatory visual population activity, and more constrained, stable early visual trajectories predicted faster responses. These population dynamics were associated with faster evidence accumulation, lower decision thresholds, and shorter non-decision time. Together, these findings suggest that associative memory facilitates perceptual decisions by shaping the content and dynamics of anticipatory neural activity, with partially dissociable contributions from visual processing and motor preparation.

**Significance statement:** Learning which events predict others helps us make perceptual decisions, but how the brain converts these associations into behavioral advantages remains unclear. We show that learned predictions preactivate expected visual information and prepare motor responses before target onset, with little evidence for enhanced visual representations afterward. Post-target visual processing and anticipatory motor preparation made partially dissociable contributions to response speed and were linked to different decision processes. Prediction also constrained the evolution of visual population activity before target onset, with shorter, more stable trajectories predicting faster responses. These findings connect associative memory to both the content and dynamics of anticipatory neural activity, showing how experience organizes the brain for perception and action.

## Introduction

Prior knowledge acquired through cue–target associative learning shapes processing across sensory, motor, and cognitive systems, enabling more efficient perception and integration of incoming information. A large body of work shows that contextual knowledge facilitates signal detection, whether through learned visual-spatial regularities linking cue and target locations^1–3^ or through regularities in which a sound object (i.e., cue) predicts when and where an auditory target will occur^4–9^. Contextual cueing provides one example of how such learned associations facilitate target processing. It has been proposed to reflect an instance-based form of automaticity, whereby memory traces from previous encounters are rapidly retrieved to guide attention and behavior^10^. However, these behavioral benefits alone do not establish whether learned associations prepare sensory and motor states before target onset, enhance sensory processing after the target appears, or both. It also remains unclear how these sensory and motor changes relate to distinct components of perceptual decision-making.

Building on this broader view of *memory-guided attention*, evidence from repetition priming suggests that prior experience can facilitate behavior through both perceptual and response-related mechanisms. Schacter et al.^11^ reviewed findings showing that priming is sensitive not only to changes in stimulus properties but also to changes in the response or decision associated with a stimulus, suggesting that perceptual representations and learned stimulus–response mappings may make partially dissociable contributions to behavioral facilitation. These response-specific priming effects may partly reflect the retrieval of learned stimulus–response associations, suggesting that associative memory influences not only perceptual processing but also downstream processes involved in response selection and action preparation. Consistent with this possibility, motor and premotor activity has been implicated in visual associative learning in macaques^12^ and in human decision-making^13^. More directly, Fischer et al.^5^ demonstrated that long-term memory for associations between everyday sound clips (cues) and lateralized tones (targets) engages the response system before target onset. Together, these findings suggest that memory-guided facilitation may comprise distinguishable sensory and response-related components.

Bayesian and predictive-processing accounts propose that prior experience and associative memories generate expectations about forthcoming events, shaping task-relevant neural activity before and during sensory processing and thereby facilitating the processing of incoming targets^14–16^. One possibility is that predictive cues activate the expected sensory representation and prepare the associated motor response before target onset, allowing the relevant neural states to be reached earlier^5,17,18^. Beyond determining which information is represented, prediction may also alter how population activity evolves over time by reshaping the dynamics and transitions among anticipatory neural states^18^. Predictive knowledge may act by preparing task-relevant sensory and motor states in advance^5,17^, by improving the quality of target processing once sensory input arrives^19–22^, or through coordinated changes in the dynamics linking perception to action. Distinguishing these possibilities is essential for understanding how learned associations enhance perceptual decision-making.

Here, we designed an associative learning paradigm using lateralized manual responses (Figure 1). Neuromagnetic brain activity was measured while participants classified visual objects as living or non-living after auditory or visual cues that either predicted the upcoming target category or provided no predictive information. Combining time-resolved representational analyses with neural-state geometry and hierarchical drift-diffusion modelling (HDDM), we tracked how prediction shaped sensory and motor activity from anticipation through decision formation and response execution. We tested whether predictive cues engage anticipatory sensory and motor preparation (anticipatory preparation hypothesis, H1), enhance post-target sensory representations (post-target enhancement hypothesis, H2), or both (Fig. 1E–G). We further examined whether visual processing and motor preparation made partially dissociable behavioral contributions and were associated with distinct latent components of perceptual decision-making. Finally, we tested whether prediction constrained visual population trajectories before and after target onset and whether these dynamics were related to motor preparation and subsequent behavior.

**Figure 1.**
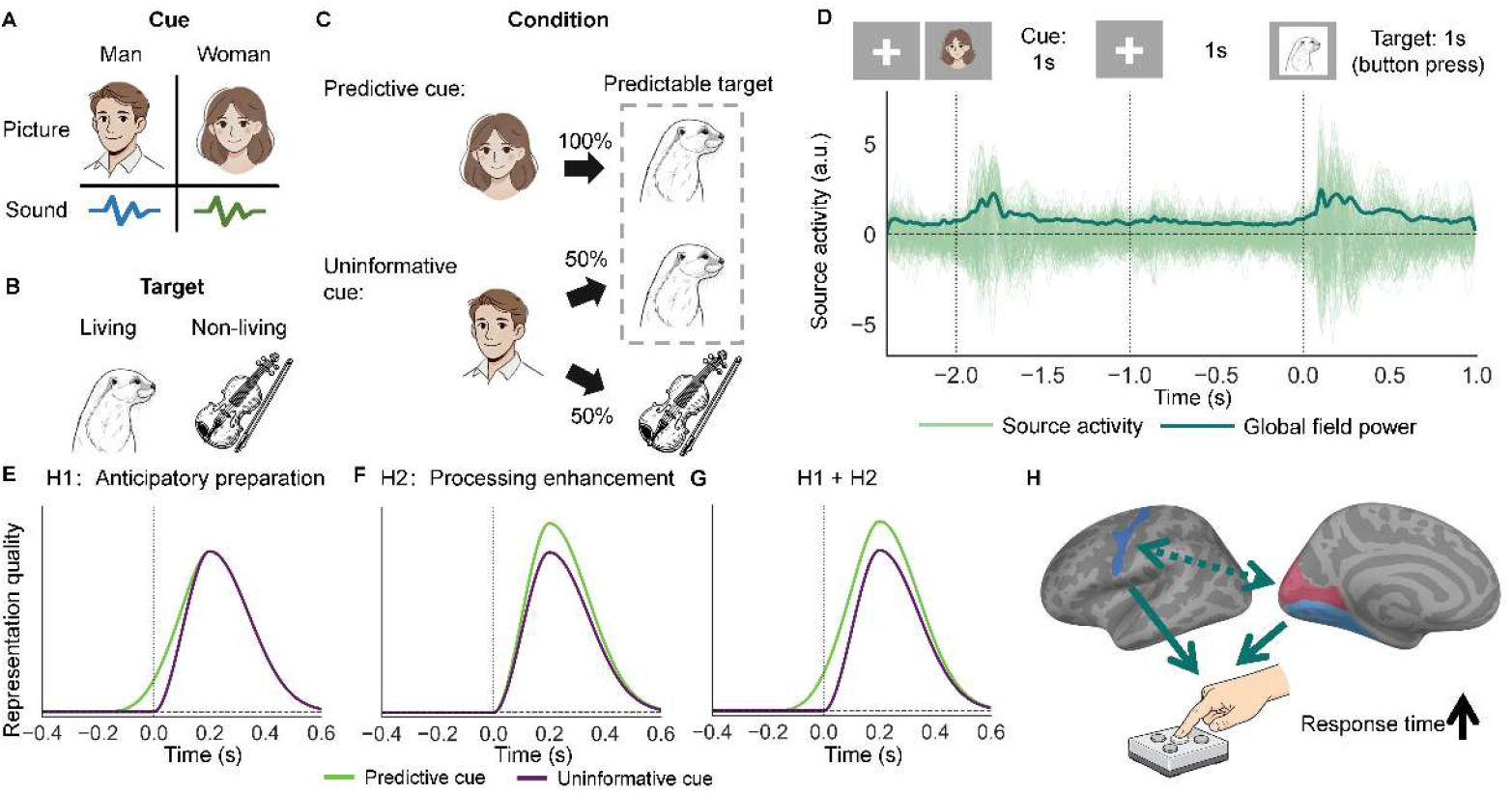
Experimental paradigm and hypothesized mechanisms of associative prediction. **A,** Auditory and visual cues depicted males or females. **B,** Target images depicted living or non-living objects, which participants categorized using left- and right-hand button presses, respectively. **C,** Within each participant, one cue type was predictive of a single target category (100%), whereas the other was uninformative and was followed equally often by living and non-living targets (50% each). **D,** Trial sequence. A 1-s auditory or visual cue was followed by a 1-s fixation interval and a 1-s target presentation, during which participants made the living/non-living judgment. Source-level activity is shown schematically across the trial, with target onset defined as time 0. **E–G,** Schematic predictions for two non-mutually exclusive mechanisms: Anticipatory preparation hypothesis (H1), predictive cues are expected to evoke target-related activity before target onset, leading to an earlier emergence of target-related representations (**E**); Processing enhancement hypothesis (H2), prediction improves the quality of target representations after stimulus onset without pre-target activation (**F**). Both mechanisms may operate concurrently (**G**). **H,** Conceptual framework in which associative prediction can facilitate behavioral responses through anticipatory sensory and motor processes and their coordination.

Analyses of sensory and motor-related brain activity revealed that predictive cues brought target-specific visual representations and response-specific motor states online before the target appeared. Beyond representational content, predictive cues also shaped how visual population activity evolved over time, resulting in more constrained and stable anticipatory trajectories. These neural effects were behaviorally consequential yet computationally distinct: visual representations were linked primarily to evidence accumulation, whereas motor preparation was linked to processes outside accumulation, and anticipatory visual population dynamics captured a broader state of behavioral readiness. Together, these findings support a multilevel mechanism in which associative predictions constrain how visual population activity patterns evolve over time before target onset, establishing a preparatory state in which target-specific sensory representations and response-specific motor activity make distinct contributions to subsequent decision-making.

## Results

### Predictive cues sped responses to predictable targets

During MEG recording, participants performed an association-learning task in which they classified each visual picture (i.e., target) as a living or non-living object as quickly and accurately as possible, using a left-hand response for living objects and a right-hand response for non-living objects. Each picture was preceded by either a predictive cue or an uninformative cue (Fig. 1A-C; see methods for details).

We first tested whether predictive cues facilitated behavioral performance. Log-transformed response times were analyzed with a linear mixed-effects model, with cue predictability as a fixed effect and participant as a random intercept. Relative to predictable-cue trials, uninformative-cue trials were associated with longer response times (Fig 2A, mean response time: 519 ms vs 532 ms, β = 0.028, t(5793) = 5.12, p < 0.001), whereas response-time variability did not differ between the two conditions (Fig 2B, p_permutation_ = 0.265). Notably, this behavioral facilitation occurred despite limited explicit awareness of these associations. When asked after the experiment which cue was predictive, only 5 of 33 participants correctly identified it; the remaining participants either reported not knowing or selected the incorrect cue. This finding suggests that predictive cues facilitated performance largely without explicit awareness of the underlying association.

**Figure 2.**
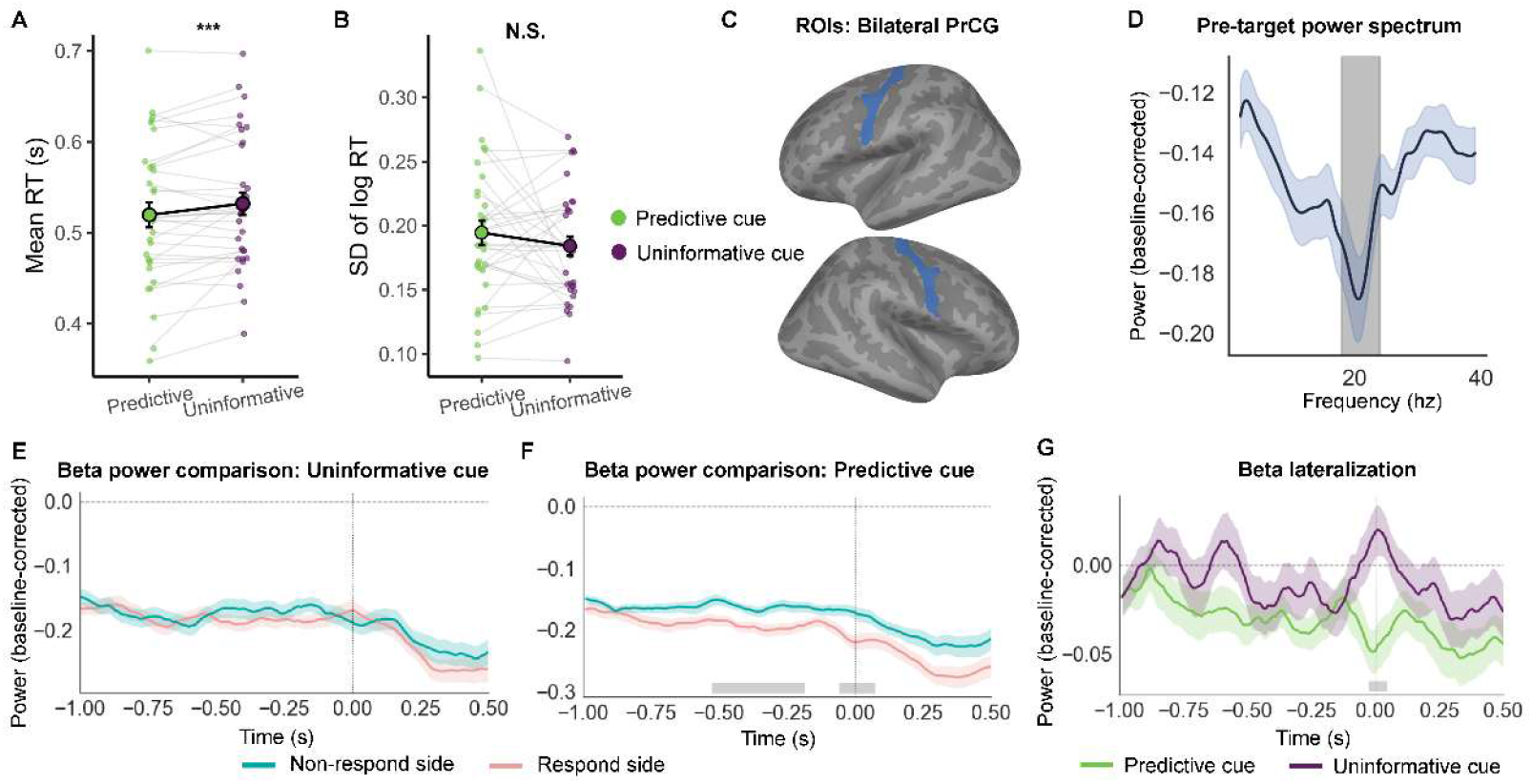
Behavioral facilitation and anticipatory response-specific motor beta suppression following predictive cues. **A,** Mean response times for predictive-target trials following predictive and uninformative cues. **B,** Within-participant variability in response time across cue conditions. Small points and connecting lines indicate individual participants, and large symbols indicate group means. **C,** Bilateral motor ROIs used for the motor beta analysis. **D,** Group-average pre-target power spectrum, showing maximal beta suppression around 21 Hz. The gray-shaded region indicates the 18–24 Hz beta range used for subsequent analyses. **E, F,** Time courses of beta power in response-related and non-response-related motor regions following uninformative (**E**) and predictive (**F**) cues. **G,** Beta lateralization with more negative values indicating stronger response-specific beta suppression. Shaded envelopes indicate SEM. Gray bars indicate significant temporal clusters. Time 0 denotes target onset. PrCG, precentral gyrus; N.S., not significant; ROIs, regions of interest; SEM, standard error of the mean.

### Predictive cues elicited anticipatory response-specific beta suppression

We next asked whether the behavioral benefit of predictive cues was accompanied by motor preparation. We focused on beta activity because its suppression over the sensorimotor cortex is a well-established marker of motor preparation^23,24^. Beta activity was quantified using source-level time-frequency representations extracted from bilateral precentral regions of interest (ROIs) (Fig. 2C). Because the group-level power spectrum showed maximal beta suppression at 21 Hz during the cue-target interval, beta power was averaged within a 6-Hz window centered on this peak, corresponding to 18-24 Hz (Fig. 2D; see methods for details).

In the uninformative-cue condition, beta power did not differ between the response-hand and non-response-hand motor regions, providing little evidence for response-specific motor preparation (Fig. 2E). In contrast, predictive cues were associated with greater beta suppression in the precentral gyrus contralateral to the response hand. This lateralized suppression was significant from −0.53 to −0.17 s before target onset (Fig. 2F, p_cluster_ = 0.001). A second cluster was observed around target onset, from −0.03 to 0.08 s (Fig. 2F, p_cluster_ = 0.009), indicating that response-specific beta suppression persisted into the earliest stage of target processing.

We then compared response-specific motor preparation using a beta lateralization index, computed as the difference in beta power between the response-hand and non-response-hand motor regions. More negative values indicated stronger suppression in the response-hand motor region. This index was significantly more negative after predictive cues than after uninformative cues immediately before target onset, between −0.03 and 0.05 s (Fig. 2G, p_cluster_ = 0.029). Thus, predictive cues enhanced the lateralized motor state just before the target appeared.

To test whether the cue-related beta-suppression effect extended beyond the motor ROIs, we repeated the analysis in the inferior frontal gyrus, opercular part (Fig. S1A-C), and the superior parietal gyrus (Fig. S1D-F). These regions were selected as anterior and posterior control regions relative to the precentral gyrus to assess the spatial specificity of the motor beta-suppression effect. Neither region showed a significant condition-related difference in beta suppression.

### Predictive cues preactivated target-specific visual representations before target presentation

We next asked whether predictive cues altered the timing and strength of neural representations of living and non-living object categories in early visual cortex (EVC) and ventral visual cortex (VVC) (Fig. 3A). The pattern specificity index (PSI) was computed as the difference between similarity to the target-matched template and to the alternative-target template, such that a PSI above zero indicated target-specific representation specificity^25,26^ (Fig. 3D; for details, see methods).

**Figure 3.**
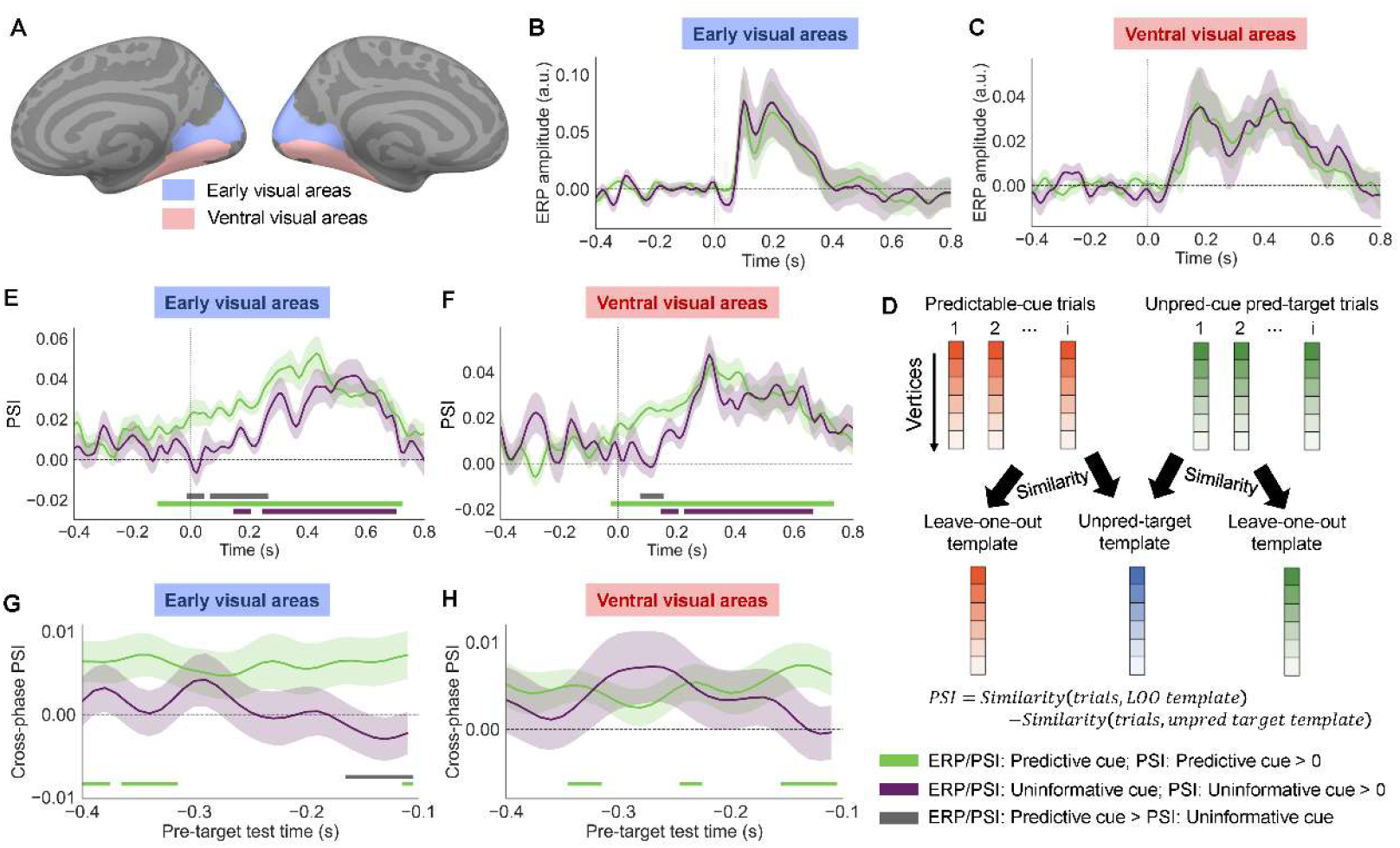
Predictive cues activated target-specific representations in visual cortex, prior to target onset. **A,** EVC and VVC regions used for source-level analyses. **B, C,** Source ERP time courses in EVC (**B**) and VVC (**C**) following predictive and uninformative cues. **D,** Schematic illustration of the PSI analysis used to quantify target-specific neural representations. **E, F,** Time-resolved PSI in EVC (**E**) and VVC (**F**). Following uninformative cues, target-specific representations emerged only after target onset, whereas predictive cues elicited target-specific activity earlier, extending into the pre-target period in EVC and around target onset in VVC. **G, H,** Cross-phase PSI in EVC (**G**) and VVC (**H**). Predictive cues elicited pre-target target-specific patterns in both visual regions, with a stronger predictive-cue effect in EVC immediately before target onset. Green and purple bars indicate intervals with above-zero PSI following predictive and uninformative cues, respectively; gray bars indicate intervals showing a significant difference between cue conditions. Shaded envelopes indicate SEM. Time 0 denotes target onset. EVC, early visual cortex; VVC, ventral visual cortex; ERP, event-related potential; PSI, pattern specificity index; a.u., arbitrary units; SEM, standard error of the mean.

We first compared source-space evoked responses and found no differences between the predictive and uninformative conditions from −0.4 to 0.8 s (Fig. 3B, C). As shown in Figure 3E, in EVC, uninformative cues elicited target-specific representations only after the target appeared, with significant above-zero PSI from 0.15 to 0.20 s (p_cluster_ = 0.019) and from 0.25 to 0.70 s (p_cluster_ < 0.001).

Strikingly, following predictive cues, significant target-specific representations were present before target onset and remained significant throughout the post-target period, from −0.11 to 0.72 s (p_cluster_ < 0.001). This pre-target activity indicates that predictive cues reinstated visual information about the expected target category before the visual stimulus was presented.

VVC showed a similar but slightly later pattern (Fig. 3F). Following uninformative cues, target-specific representations emerged after target onset, from 0.15 to 0.20 s (p_cluster_ = 0.031) and from 0.23 to 0.66 s (p_cluster_ < 0.001). Following predictive cues, PSI became significant around target onset and remained elevated from −0.02 to 0.73 s (p_cluster_ < 0.001), suggesting that predictive cues also facilitated target-specific processing along the ventral visual pathway.

Finally, we tested whether these anticipatory representations resembled those evoked by the subsequent target. Cross-phase PSI was averaged over the post-target windows that showed significant differences between the two conditions in the preceding analysis. Predictive cues produced significant pre-target PSI in both EVC (−0.40 to −0.38 s, −0.36 to −0.32 s, and −0.11 s) and VVC (−0.34 to −0.32 s, −0.24 to −0.23 s, and −0.15 to −0.11 s), whereas no significant PSI was observed following uninformative cues (Fig. 3G, H). Critically, cross-phase PSI was greater for predictive than for uninformative cues in EVC from −0.16 to −0.11 s (Fig. 3G, *p*_cluster_ = 0.005). This finding suggests that predictive cues activated target-evoked visual representations before target onset.

### Pre-target motor preparation and post-target visual specificity each accounted for unique variance in response speed

We next asked whether trial-by-trial fluctuations in the neural signals identified above predicted behavioral performance, assessed using trial-resolved neural–behavioral coupling analysis. This analysis was restricted to predictable-target trials, and cue predictability was included as a covariate in all models. Log-transformed response time was modeled using linear mixed-effects models^27^, with neural activity as a fixed effect and participant as a random intercept. Neural predictors were z-scored within participants before model fitting.

Consistent with a role in anticipatory motor preparation, beta lateralization in the pre-target window from −0.53 to −0.17 s significantly predicted response time (Fig. 4A, β = 0.0049, t(5416) = 2.31, p = 0.021), indicating that stronger pre-target beta suppression was associated with faster responses. In contrast, beta lateralization in the peri-target window from −0.03 to 0.08 s was not significantly associated with response time (β = 0.0030, t(5416) = 1.43, p = 0.153). We then tested whether target-specific visual representations were also related to behavior. PSI in the early post-target window significantly predicted response time, with stronger target-specific representations associated with faster responses (Fig. 4B, β = −0.0050, t(5416) = −2.36, p = 0.018), whereas PSI in the earlier window was not significantly related to response time (β = −0.0013, t(5416) = −0.61, p = 0.541).

**Figure 4.**
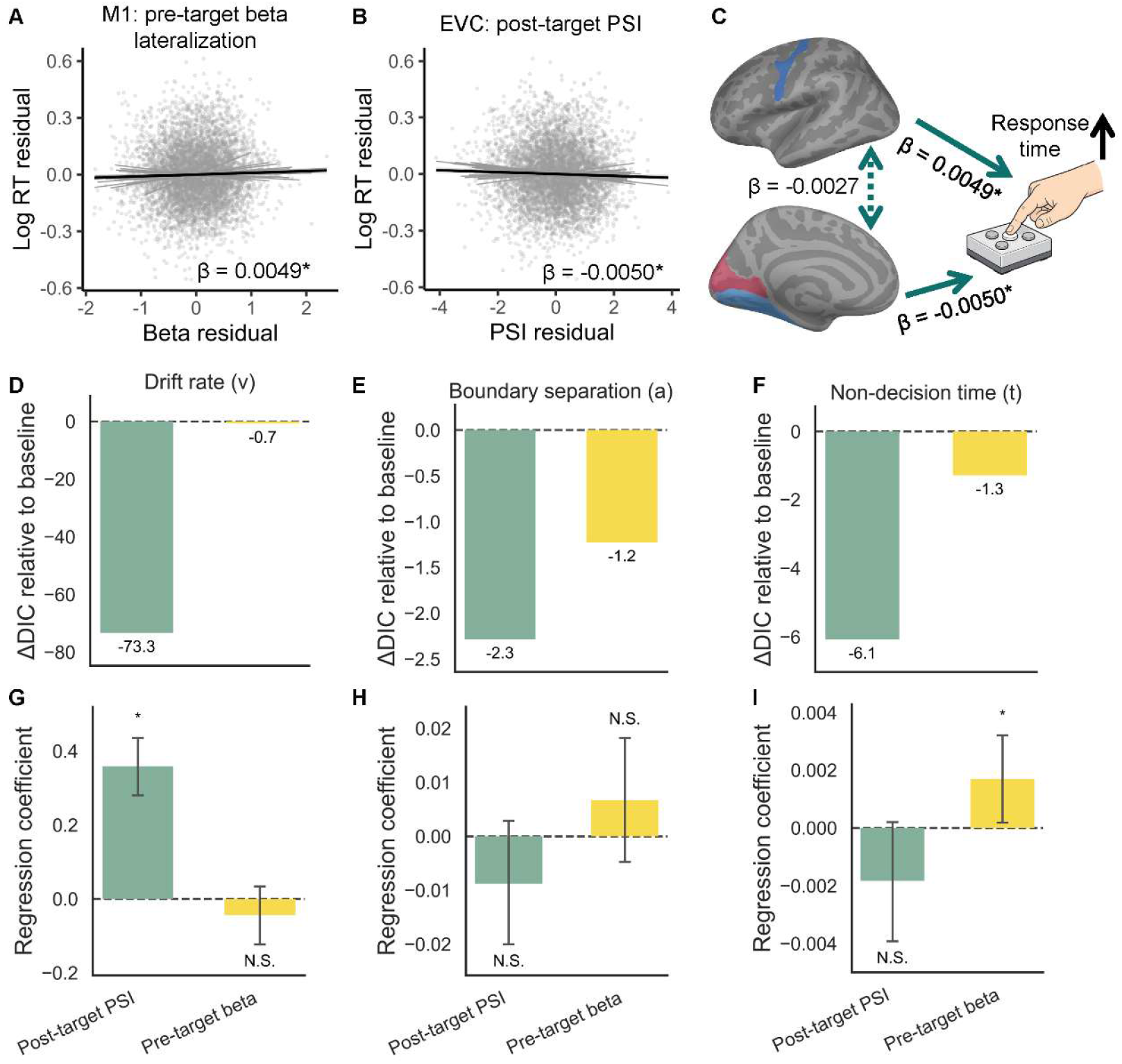
Pre-target motor preparation and post-target visual specificity independently predicted response speed and mapped onto distinct decision components. A, B, Trial-resolved neural–behavioral associations between response time and pre-target motor beta lateralization (A) or post-target PSI in EVC (B). Stronger pre-target response-specific beta suppression and greater post-target visual specificity were each associated with faster responses. **C,** Summary of the joint model including both neural measures, showing that pre-target motor beta lateralization and post-target PSI accounted for unique variance in response speed. The dashed connection denotes a nonsignificant correlation. **D–F,** Changes in model fit relative to the behavioral baseline HDDM for models in which post-target PSI or pre-target motor beta lateralization modulated drift rate (**D**), boundary separation (**E**), or non-decision time (**F**). More negative ΔDIC values indicate greater improvement in model fit. **G–I,** Posterior regression coefficients relating the two neural measures to drift rate (**G**), boundary separation (**H**), and non-decision time (**I**). Post-target PSI was selectively associated with faster evidence accumulation, whereas stronger pre-target response-specific beta suppression was associated with shorter non-decision time; neither neural measure was reliably associated with boundary separation. Error bars indicate 95% credible intervals. M1, precentral gyrus; EVC, early visual cortex; PSI, pattern specificity index; RT, response time; HDDM, hierarchical drift-diffusion modeling; DIC, deviance information criterion; N.S., not significant.

Finally, when pre-target motor beta lateralization and post-target PSI were entered into the same model, both effects remained significant. Stronger pre-target beta suppression predicted faster responses (β = 0.0049, t(5415) = 2.31, p = 0.021), and stronger post-target PSI also predicted faster responses (β = −0.0050, t(5415) = −2.36, p = 0.018). These results indicate that anticipatory motor beta suppression and early target-specific visual processing accounted for unique variance in response speed beyond the overall effect of cue predictability (Fig. 4C).

Together, these findings from the analysis of neural activity in the motor and visual cortices support the anticipatory preparation hypothesis, showing that predictive cues engage response-specific motor preparation and target-specific visual representations before target onset, but provide little evidence of post-target enhancement.

### Motor preparation and visual specificity mapped onto distinct decision components

We then tested whether anticipatory motor preparation and target-specific visual processing mapped onto distinct components of perceptual decision-making. We fitted HDDM to trial-by-trial choices and response times, using motor beta lateralization or PSI as a neural predictor of drift rate (v), boundary separation (a), and non-decision time (t), which respectively capture the rate of evidence accumulation, response caution, and processing outside evidence accumulation, including sensory encoding and motor execution^28^.

Pre-target motor beta lateralization was associated with non-decision time (posterior mean = 0.0017, 95% CI [0.0002, 0.0032]; DIC = −943.60): stronger response-specific beta suppression was associated with shorter processing time outside evidence accumulation.

Post-target PSI quantified the relative similarity of visual activity to the target-matched versus alternative-target template. Higher trial-level PSI therefore indicated more target-specific visual representations. Higher PSI was associated with faster evidence accumulation toward the target-consistent response (drift-rate coefficient: posterior mean = 0.359, 95% CI [0.280, 0.436]). Including PSI as a predictor of drift rate substantially improved model fit relative to the behavioral baseline (DIC = −1015.65 vs. −942.31).

Motor beta lateralization was not reliably associated with drift rate, whereas PSI was not reliably associated with non-decision time. Neither measure was reliably associated with boundary separation. Thus, visual and motor signals mapped onto distinct components of the decision process: target-specific visual activity tracked the efficiency of evidence accumulation, whereas anticipatory motor preparation was linked to processes outside the accumulation stage.

### Predictive cues constrained anticipatory population dynamics in the visual cortex

The preceding analyses showed that predictive cues activated target-specific visual representations before target onset. However, representational analyses establish what information is present at a given time, but not how the distributed population state evolves during anticipation. This distinction is important because similar representational content can arise under different temporal dynamics. We therefore used principal component analysis (PCA) to represent EVC and VVC activity in a low-dimensional state space. Each axis represented a principal component of activity across vertices within an ROI, and each point represented the population activity pattern at a given time. Successive points formed a trajectory, allowing us to test whether prediction constrained the temporal evolution of population activity. Changes between consecutive states were quantified as Euclidean distances between adjacent trajectory points. We focused on trajectory dynamics from −0.40 to −0.10 s, quantified by the total distance travelled through state space and the variability of consecutive state transitions.

In both EVC and VVC, neural population activity patterns showed significantly less cumulative change over time following predictive than uninformative cues (Fig. 5B, E; EVC: mean difference = 3.14, p_permutation_ = 0.003; VVC: mean difference = 2.39, p_permutation_ = 0.012), indicating less extensive state-space evolution during anticipation under prediction. In EVC, this effect was accompanied by significantly reduced variability in consecutive step distances (Fig. 5C; mean difference = 0.043, p_permutation_ = 0.021), signifying more stable moment-to-moment population dynamics. VVC showed the same numerical tendency, although the reduction in step-distance variability did not reach significance (Fig. 5F; mean difference = 0.027, p_permutation_ = 0.078). We also repeated the trajectory analyses in the post-target interval from 0.10 to 0.40 s. Neither total path length nor step-distance variability differed significantly between predictive and uninformative conditions in EVC or VVC. Thus, effects of prediction on these trajectory measures were detected during anticipation, but not in the tested post-target interval.

**Figure 5.**
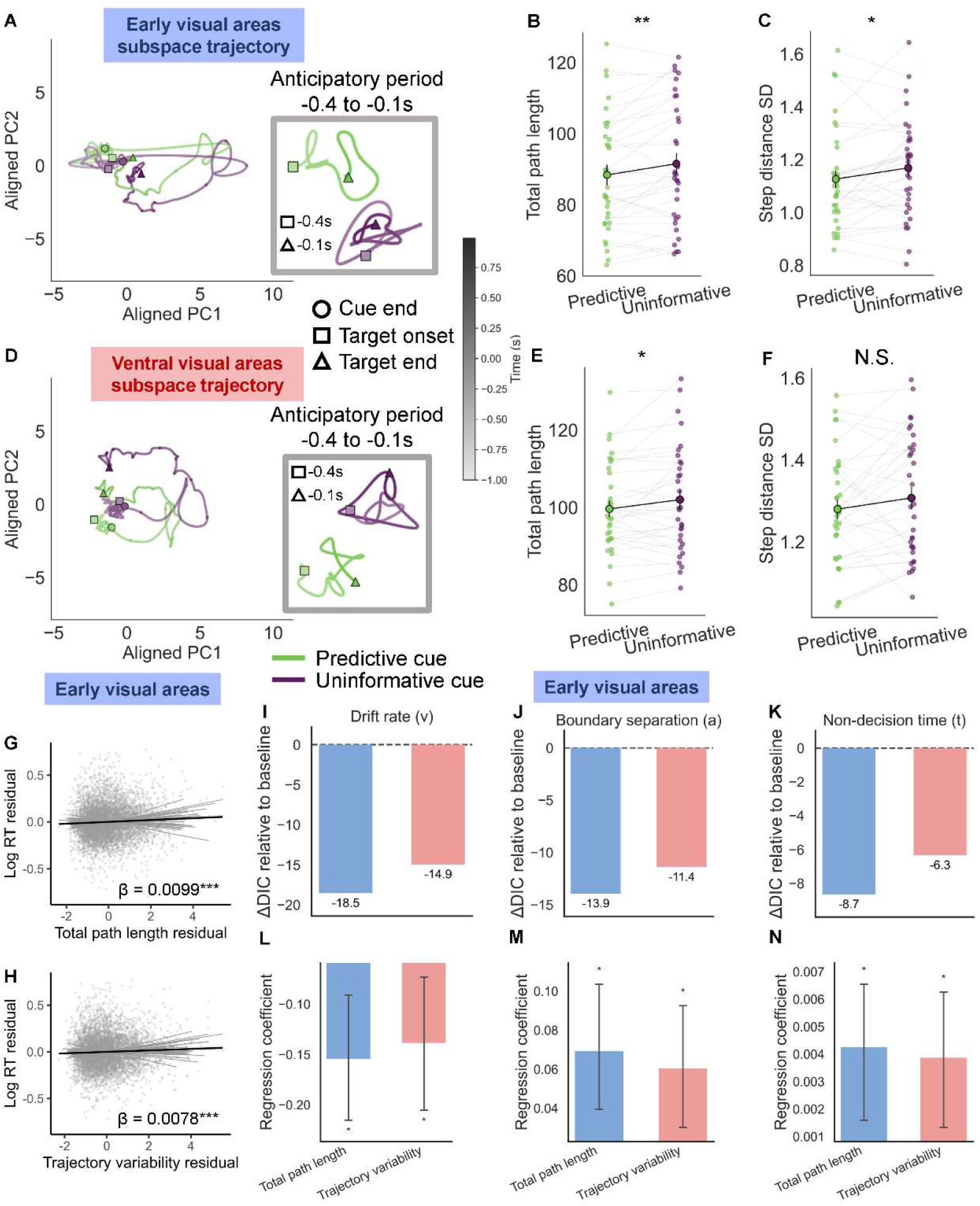
Predictive cues constrained anticipatory visual population dynamics, which predicted response speed and multiple components of decision processing. A, D, Population trajectories through a low-dimensional neural state space in EVC (**A**) and VVC (**D**) following predictive and uninformative cues. Insets show trajectories during the anticipatory period from −0.40 to −0.10 s. Symbols mark cue offset, target onset, and target offset. **B, C,** Anticipatory trajectory measures in EVC. Predictive cues were associated with shorter total path length (**B**) and lower step-distance variability (**C**), indicating less extensive and more stable moment-to-moment state-space evolution. **E, F,** Corresponding trajectory measures in VVC. Predictive cues were associated with shorter total path length (**E**), whereas step-distance variability did not differ reliably between cue conditions (**F**). **G, H,** Trial-resolved associations between response time and EVC total path length (**G**) or step-distance variability (**H**). Shorter and more stable anticipatory trajectories were associated with faster responses. **I–K,** Changes in model fit relative to the behavioral baseline HDDM for models in which EVC total path length or step-distance variability modulated drift rate (**I**), boundary separation (**J**), or non-decision time (**K**). **L–N,** Posterior regression coefficients relating EVC total path length and step-distance variability to drift rate (**L**), boundary separation (**M**), and non-decision time (**N**). More constrained and stable anticipatory EVC dynamics were associated with faster evidence accumulation, lower decision thresholds, and shorter non-decision time. Error bars indicate 95% credible intervals. EVC, early visual cortex; VVC, ventral visual cortex; PC, principal component; RT, response time; HDDM, hierarchical drift-diffusion modeling; DIC, deviance information criterion; SD, standard deviation; N.S., not significant.

Because predictive cues also elicited anticipatory motor beta suppression, we tested whether the trajectory changes observed in the visual cortex extended to the precentral gyrus. We therefore applied the same trajectory analysis to pre-target activity in the precentral gyrus. Neither total path length (mean difference = 0.93, p_permutation_ = 0.116) nor step-distance variability (mean difference = −0.002, p_permutation_ = 0.572) differed significantly between predictive and uninformative conditions (Fig. S2). Thus, the motor beta effect was not accompanied by detectable changes in these M1 trajectory measures.

Together, these findings extend prior evidence for target-specific preactivation by revealing a complementary effect of prediction on population dynamics. Predictive cues not only reinstated information about the expected target but also constrained the pre-target evolution of visual population activity, producing smaller cumulative changes in population activity patterns in both EVC and VVC and more stable moment-to-moment dynamics in EVC. The absence of comparable effects in M1 further suggests that this dynamical stabilization was preferentially expressed within the visual cortex rather than reflecting a global consequence of predictive cueing. Thus, predictions based on associative memory shaped anticipatory visual processing at two complementary levels: they activated the content of the expected representation while organizing visual population activity into a more constrained and stable dynamical regime before target onset.

### Anticipatory EVC population dynamics predicted response speed and multiple components of decision processing

Using our trial-resolved neural–behavioral coupling analysis, we examined whether trial-to-trial fluctuations in anticipatory population dynamics were related to subsequent behavioral performance. EVC total path length significantly predicted response time (Fig. 5G, β = 0.0099, *t*(5583) = 4.23, *p* < 0.001), indicating that less extensive state-space movement was associated with faster responses. Similarly, EVC step-distance variability significantly predicted response time (Fig. 5H, β = 0.0078, *t*(5583) = 3.35, *p* < 0.001), with more stable moment-to-moment population dynamics associated with faster responses. In contrast, neither total path length nor step-distance variability in VVC was significantly associated with response time (total path length: β = 0.0018, *t*(5583) = 0.76, *p* = 0.448; step-distance variability: β = 0.0020, *t*(5583) = 0.86, *p* = 0.392).

We further used HDDM to determine which components of decision processing were associated with these EVC dynamics. Greater total path length was associated with a lower drift rate (Fig. 5I, L, posterior mean = −0.155, 95% CI [−0.215, −0.091], ΔDIC = −18.52), greater boundary separation (Fig. 5J, M, posterior mean = 0.069, 95% CI [0.039, 0.104], ΔDIC = −13.94), and longer non-decision time (Fig. 5K, N, posterior mean = 0.0043, 95% CI [0.0016, 0.0065], ΔDIC = −8.65). Greater step-distance variability showed the same pattern, with a lower drift rate (Fig. 5I, L, posterior mean = −0.139, 95% CI [−0.205, −0.073], ΔDIC = −14.93), greater boundary separation (Fig. 5J, M, posterior mean = 0.060, 95% CI [0.030, 0.093], ΔDIC = −11.39), and longer non-decision time (Fig. 5K, N, posterior mean = 0.0039, 95% CI [0.0013, 0.0063], ΔDIC = −6.34). All six neural-regressor models therefore improved model fit relative to the baseline HDDM, with the largest improvements observed in models in which EVC population dynamics modulated drift rate.

Notably, associations between EVC population dynamics and response time were more robust than those observed for pre-target motor beta lateralization or post-target visual PSI. Moreover, these signals showed more parameter-specific relationships with decision processing, with EVC trajectory measures associated with drift rate, boundary separation, and non-decision time. This broader pattern suggests that anticipatory population dynamics may capture a more global preparatory state that influences multiple components of subsequent response formation.

Together, these results indicate that trials with more constrained and stable EVC population dynamics were followed by faster responses, faster evidence accumulation, lower decision thresholds, and shorter non-decision processes. These findings are consistent with the anticipatory preparation hypothesis, showing that the organization of visual population activity before target onset was associated with subsequent response speed and decision parameters.

### Visual and motor anticipatory signals were coupled yet independently related to behavior

We next examined whether anticipatory visual population dynamics were coupled with motor preparation. EVC trajectory variability significantly predicted pre-target beta lateralization (β = 0.015, *t*(5584) = 2.19, *p* = 0.029), with lower trajectory variability linked to stronger beta suppression in the response hand (Fig. 6A). In contrast, EVC total path length and both VVC trajectory measures were not significantly associated with beta lateralization.

**Figure 6.**
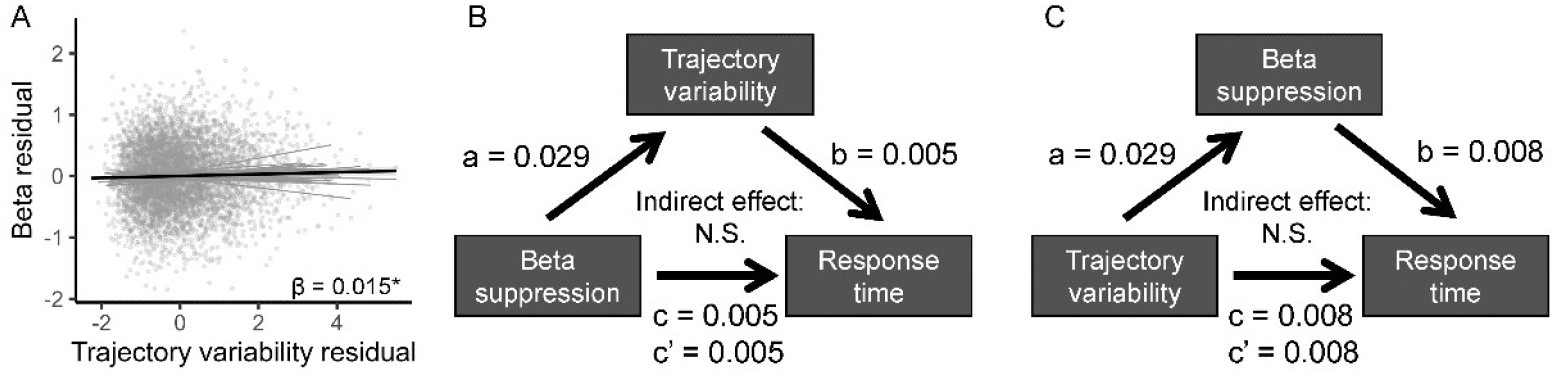
Visual and motor anticipatory signals were coupled but independently related to behavior. **A,** Trial-resolved association between EVC trajectory variability and pre-target motor beta lateralization. Lower trajectory variability was associated with stronger response-specific beta suppression. **B, C,** Bidirectional multilevel mediation analyses testing whether EVC trajectory variability mediated the relationship between motor beta lateralization and response time (**B**), or whether motor beta lateralization mediated the relationship between trajectory variability and response time (**C**). Neither indirect effect was reliable. Paths *a*, *b*, *c*, and *c′* denote mediation coefficients. EVC, early visual cortex; N.S., not significant.

We then tested whether this visual–motor coupling mediated their relationships with behavior. Multilevel mediation analyses provided no evidence of mediation in either direction. When EVC trajectory variability was modeled as mediating the relationship between motor beta lateralization and response time, the indirect effect was likewise unreliable (Fig. 6B, indirect effect = 0.00006, 95% CI [−0.00027, 0.00063]). Likewise when motor beta lateralization was modeled as mediating the relationship between EVC trajectory variability and response time, the indirect effect was not reliable (Fig. 6C, indirect effect = 0.00013, 95% CI [−0.00017, 0.00046]). Thus, although anticipatory visual population dynamics covaried with motor preparation, their associations with behavior were not explained by a simple mediated pathway, suggesting that visual-state stabilization and motor preparation made partially dissociable contributions to subsequent responses.

## Discussion

Memory shapes how we allocate attention and respond to incoming sensory stimuli. Here, we show that associative memory facilitates perceptual decisions by coordinating neural processing before sensory evidence arrives. Predictive cues accelerated object categorization despite limited explicit awareness of the cue–target associations, while simultaneously activating target-specific visual representations, engaging response-specific motor preparation, and constraining the evolution of anticipatory activity in the visual cortex. Critically, these anticipatory dynamics were behaviorally consequential: more constrained and stable EVC population trajectories predicted faster responses and were associated with multiple components of subsequent decision formation. By contrast, target-specific visual representations and motor preparation showed more selective computational relationships, with post-target visual specificity linked primarily to evidence accumulation and anticipatory motor activity linked to non-decision processing. Although visual population stability covaried with motor preparation, their behavioral effects were not explained by a simple serial mediation pathway. Together, these findings suggest that associative memory facilitates perceptual decision-making performance by guiding attention toward expected target categories, shaping anticipatory visual population dynamics, and engaging sensory and motor processes that make partly dissociable contributions to subsequent decision-making.

Our findings extend classic accounts of contextual cueing and memory-guided attention^1–3,6,7,9,29,30^. Classic contextual-cueing studies have shown that repeated spatial configurations enable observers to implicitly learn predictive associations, which in turn guide attention more efficiently toward task-relevant locations^1–3^. According to instance-based accounts of automaticity^10^, these benefits arise because memory traces from prior encounters are rapidly reactivated, allowing learned context to bias processing on subsequent encounters. Our results are consistent with this general framework, particularly given that predictive cues facilitated behavior despite limited explicit awareness of the cue–target associations. Related work on repetition priming has proposed that experience can also improve processing efficiency through enhanced neural synchronization^31,32^. Here, we show that learned cue–target associations shaped visual processing before the target appeared, both by reinstating target-specific information and by constraining the evolution of anticipatory population activity. Thus, memory-guided facilitation may operate not only by directing attention to behaviorally relevant information, but also by configuring the sensory system into a more stable, task-ready state before sensory evidence becomes available.

One component of this anticipatory state was the activation of target-specific visual representations. Predictive cues elicited target-specific activity before target onset, whereas comparable representations following uninformative cues emerged only after the target appeared. This shift was particularly evident in EVC, where prediction advanced target-related information by approximately 270 ms. Cross-phase analyses further showed that pre-target activity resembled post-target target-specific patterns, indicating reinstatement of the expected sensory representation. These findings are consistent with predictive-coding accounts in which prior knowledge generates sensory templates before bottom-up input^14,15,17^, and extend them by showing that learned cue–target associations can activate behaviorally relevant object-category information before the stimulus appears. This pattern supports our first hypothesis that associative predictions facilitate perception by bringing expected sensory representations online earlier. By contrast, we found no evidence that prediction further enhanced the quality of target-specific representations after stimulus onset, providing little support for the alternative processing-enhancement account.

Prediction, however, altered not only which information was represented during anticipation, but also how visual population activity evolved over time. Population-dynamics frameworks emphasize that neural computation depends not only on representational content at individual time points but also on the trajectories by which population states evolve^33–35^. Previous work further suggests that prestimulus population states can constrain subsequent perceptual processing^36,37^, and that expectation can reshape ongoing population dynamics before stimulus arrival, facilitating subsequent sensory coding^18^. Consistent with these accounts, we found that associative memory altered prestimulus population dynamics, but in the form of more constrained activity in visual cortices: predictive cues reduced the extent of state-space evolution in both EVC and VVC and reduced moment-to-moment variability in EVC. Importantly, these dynamics were behaviorally relevant: in EVC, shorter and more stable trajectories predicted faster responses and were associated with faster evidence accumulation, lower decision thresholds, and shorter non-decision time. Thus, associative memory may facilitate perception not only by activating expected sensory content, but also by organizing visual population activity into a constrained anticipatory state that supports more efficient subsequent decision processing.

However, emerging evidence suggests that memory-guided behavior is not solely a product of attentional facilitation^5,6,38^. A related distinction has long been recognized in the repetition-priming literature, where behavioral facilitation can reflect not only changes in perceptual representations but also learned stimulus–response associations^38,39^. Such response-learning accounts raise the possibility that retrieving a familiar association may prepare not only what is likely to be perceived, but also how the organism is likely to respond. Consistent with this view, recent work has implicated motor and premotor systems in perceptual decision-making even before overt action is required^5,12^. Fischer and colleagues^5^, for example, demonstrated using EEG that long-term associative memories can pre-activate motor programs, effectively preparing the response system in advance of stimulus onset and thereby improving task performance. Extending this work, we found that predictive cues elicited response-specific motor beta suppression before target onset, indicating anticipatory preparation of the motor effector associated with the expected response. Importantly, this motor preparation occurred alongside target-specific visual activation, suggesting that associative memory can concurrently prepare sensory and motor systems before incoming sensory evidence becomes available.

Previous neural decision-modeling studies have shown that trial-by-trial neural signals relate to distinct latent components of perceptual decision-making, with stimulus-specific sensory activity particularly linked to the accumulation of perceptual evidence^40,41^. Similarly, studies using traditional spatial-and feature-based attention paradigms have shown that attentional allocation can modulate evidence accumulation and non-decision time^42,43^. Our findings extend this framework to memory-guided perceptual decisions, showing that sensory and motor signals recruited by associative prediction make partially dissociable contributions to behavior. Trial-by-trial neural–behavioral analyses showed that pre-target motor preparation and post-target visual specificity each independently predicted faster responses, indicating that their behavioral associations were not redundant. HDDM further revealed a functional dissociation: post-target visual specificity was primarily associated with increased drift rate, indicating more efficient accumulation of sensory evidence, whereas anticipatory motor beta suppression was associated with reduced non-decision time, indicating shorter processing outside evidence accumulation. Thus, within the same memory-guided decision, target-specific sensory processing and anticipatory motor preparation were linked to distinct latent components of response formation. These findings suggest that associative memory facilitates behavior through parallel but functionally differentiated sensory and motor pathways, influencing different stages of the transformation from sensory evidence to action.

Future work could further dissociate complementary dimensions of associative prediction. Independently manipulating target identity and cue–target timing would help separate “what” from “when” predictions, particularly because the present paradigm used a fixed 1-s interval between cue offset and target onset. Likewise, independently manipulating target identity and response mapping could more directly dissociate sensory from motor prediction. Together, these designs would clarify how content, timing, and response expectations jointly shape anticipatory sensory and motor dynamics.

Our findings support a framework in which associative memory facilitates visual decisions through multiple, partially dissociable neural mechanisms. Predictive cues organize anticipatory visual population dynamics while concurrently activating target-specific sensory representations and response-specific motor states. These signals serve complementary functions: anticipatory visual dynamics capture a broader preparatory state, whereas sensory and motor signals make more selective contributions to subsequent decision-making. This integrated perspective extends traditional accounts of memory-guided attention by showing that associative memory does more than bias attentional selection; it reshapes the evolving neural state from which perceptual decisions arise.

## Methods

### Participants

Thirty-three young adults (22.58 ± 4.54 years, 18 female) participated in this study. Sample size was determined through an a priori power analysis using G*Power version 3.1.9.7^44^. This sample size was sufficient to detect a medium effect size (0.5) with 80% power using a paired, two-tailed t-test. All participants were healthy, right-handed individuals with no history of neurological disorder and with normal hearing and vision. All participants provided written informed consent, which was approved by the Baycrest Research Ethics Board (Approval No. 24-33).

### Stimuli

The stimuli comprised visual and auditory cues and picture targets (Fig. 1A). Visual cues were pictures of a male or a female. Auditory cues were 1-s /la/ syllables spoken in a male or female voice, generated in Synthesizer V Studio Basic with the pitch set to E4. Target stimuli were pictures of living and non-living items, such as animals and everyday objects (Fig. 1B). The initial target set was selected from the International Picture Naming Project^45^, which provides 520 common object drawings and 275 action drawings with naming norms. Additional target pictures were generated with Gemini 1.5 to match the visual style of the database stimuli.

### Procedure

Participants performed a cue-target semantic decision task (Fig. 1D). Each trial began with a visual or auditory cue presented for 1 s. After the cue, a fixation cross was presented for 1 s, followed by the target picture for 1 s. The inter-trial interval after target presentation varied from 1.5 to 2.0 s in 0.1 s increments.

Cue-target associations were manipulated within each participant so that one cue type predicted the upcoming target category, whereas the other was non-predictive (Fig. 1C). In the predictive condition, all targets following a male or female cue belonged to a single semantic category (living or non-living, depending on the cue). In the non-predictive condition, the cue was followed equally often by living and non-living targets. For example, in one assignment, male cues were always followed by non-living targets, whereas female cues were followed by either living or non-living targets. The assignment of predictive cue type and predicted target category was counterbalanced across participants.

Participants were instructed to judge whether the target was living or non-living as quickly and accurately as possible. No response was required to the cue. Responses were made using two buttons, with the left hand assigned to living targets and the right hand to non-living targets. The response mapping remained fixed throughout the experiment.

Each participant completed three blocks of 80 trials, except for one participant who completed 60 trials per block due to a technical issue. Within each block, male and female cues were presented equally often, with 40 trials per cue type. Within each cue type, visual and auditory cues were also presented equally often, with 20 picture cues and 20 sound cues. Thus, cue type and cue modality were balanced within each block. Target images were selected without replacement from separate living and non-living image pools. Trial order was pseudo-randomized with two constraints: no more than four consecutive trials could have the same target category, and no more than three consecutive trials could have the same cue modality.

### Behavioral analysis

Response times were log-transformed before statistical analysis^46^. Because cue modality did not interact with cue predictability, subsequent analyses collapsed across visual and auditory cue trials. To assess whether cue predictability facilitated behavioral performance, log-transformed response times were modeled using a linear mixed-effects model with cue predictability as a fixed effect and participant as a random intercept:

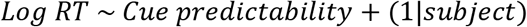

Trial-level associations were assessed using linear mixed-effects models implemented in R with the lme4 and lmerTest packages^46,47^.

Response-time variability was quantified as the within-participant standard deviation of log-transformed response times after trial-number matching between cue conditions, averaged across 1,000 resamples, and compared using a two-sided paired sign-flip permutation test with 100,000 permutations.

### MEG data acquisition and preprocessing

MEG data were recorded using a 275-channel CTF system (VSM MedTech) at a sampling rate of 1200 Hz. Continuous data were band-pass filtered between 0.1 and 120 Hz using an FIR filter. To remove artifacts, we applied independent component analysis to the sensor-level MEG data. ICA was fitted to a 1–40 Hz filtered copy of the data using the Infomax algorithm with 50 components. Components reflecting eye blinks, saccades, or cardiac artifacts were identified through visual inspection of component time courses and topographies and then removed. The cleaned data were epoched from −2.5 to 1.0 s, time-locked to the living and non-living target onsets. Epochs were baseline-corrected using the −2.30 to −2.02 s pre-cue baseline and down-sampled to 250 Hz. Epochs were then manually inspected to reject residual artifacts. MEG data preprocessing and analysis were performed using MNE-python 1.9.0^48^.

### Structural data acquisition and source localization

Structural MRI data were collected on a Siemens 3T Prisma scanner with a 64-channel head coil. T1-weighted images were acquired using the magnetization-prepared rapid acquisition gradient-echo (MPRAGE) sequence (TR = 2000 ms, TE = 2.85 ms, field of view = 256 × 240 mm, and voxel size = 0.8 × 0.8 × 0.8 mm).

Structural T1-weighted MRIs were processed with FreeSurfer to reconstruct cortical surfaces and define the source space^49^. MEG–MRI coregistration was performed individually, run by run, by aligning fiducial points to anatomical landmarks (nasion and bilateral preauricular points). A single-shell boundary-element model was generated to represent the inner-skull conductivity. Surface source spaces were constructed using an oct6 cortical grid, and individual forward solutions were computed for each experimental run. Linearly constrained minimum variance spatial filters were constructed separately for each run using empirical data covariance from −2.0 to 1.0 s and noise covariance from −2.48 to −2.02 s. Filters were built with max-power orientation, 0.05 regularization, and unit-noise-gain normalization, and were then applied to cleaned MEG epochs to obtain single-trial source time courses. Epochs with head displacements exceeding 10 mm were excluded. Head displacement was computed from the circumcenter of three localization coils for each epoch.

Source estimates were cropped from −2.4 to 1.0 s, low-pass filtered at 40 Hz, and down-sampled to 100 Hz. All source data were computed in native space and parcellated using the Destrieux cortical atlas (aparc.a2009s) for subsequent ROI-based decoding analyses^50^.

### Motor beta suppression analysis

Motor beta activity was quantified from source-level time-frequency representations in the bilateral motor cortices. For each participant, source time courses were extracted from the superior portions of the left and right precentral gyri, defined by the Destrieux G_precentral labels^50^. Within each ROI, vertex-wise source activity was summarized using the first principal component with sign correction (pca_flip). Trials were grouped by cue predictability and target category. Time-frequency power was computed using Morlet wavelets from 2 to 40 Hz in 1-Hz steps, with the number of cycles set to half the frequency. Power was baseline-corrected using a log-ratio transformation relative to the pre-cue baseline (-2.3 to -2.02 s). The beta frequency of interest was defined empirically from the group-level motor power spectrum. Power was first averaged across the −1.0 to 0 s cue-target interval, and the frequency showing the strongest beta suppression was identified at 21 Hz. We therefore defined the beta band as a 6-Hz window centered on this peak, corresponding to 18-24 Hz.

For group-level analyses, left and right motor ROIs were recoded based on the response hand. Because living targets were associated with left-hand responses and non-living targets with right-hand responses, response-related motor activity was defined as right precentral activity for living targets and left precentral activity for non-living targets. Non-response motor activity was defined as activity in the opposite hemisphere.

Motor beta suppression was quantified by averaging power in the 18-24 Hz range. Time courses were visualized from −1.0 to 0.5 s relative to target onset. Statistical testing was restricted to the −1.0 to 0.2 s interval because the fastest behavioral responses occurred at approximately 0.2 s after target onset; this restriction minimized potential contamination from overt motor execution. Response-hand and non-response-hand beta power were compared using paired-sample cluster-based permutation tests across time, with a cluster-forming threshold of p < 0.005, 2,000 permutations, and a cluster-level alpha of 0.05.

### Pattern specificity index analysis

The pattern specificity index (PSI) was quantified using a time-resolved trial-to-template similarity analysis of source-level activity. PSI indexes the extent to which the neural activity pattern at a given time point more closely resembles the representation of the current target category than the alternative target category^25,26^. Higher PSI values therefore indicate greater target-specific representational specificity. The analysis focused on two visual macro-regions of interest: EVC and VVC (Fig. 3A). EVC included bilateral S_calcarine, G_cuneus, and G_occipital_sup labels. VVC included the bilateral G_oc-temp_lat-fusifor, G_oc-temp_med-Lingual, S_oc-temp_med_and_Lingual, and S_collat_transv_ant labels. For each participant and label, single-trial vertex-wise source activity was extracted and band-pass filtered between 1 and 40 Hz. Activity was baseline-corrected relative to the pre-cue interval from −2.3 to −2.02 s.

For each label and time point, we computed the spatial correlation between each trial’s vertex-wise activity pattern and a condition-specific template pattern (Fig. 3D). Within-condition similarity was computed using a leave-one-trial-out template to avoid circularity. PSI was defined as the difference between similarity to the target-matched template and similarity to the alternative-target template. For predictive cue trials, PSI was calculated as the similarity among predictable-cue trials minus the similarity between predictable-cue trials and the uninformative cue alternative-target template. For uninformative cue trials containing the predictable target, PSI was calculated as the similarity among uninformative cue predictable-target trials minus the similarity between these trials and the uninformative cue alternative-target template. Higher PSI values, therefore, indicate stronger target-specific neural representations.

For group-level analyses, label-wise PSI time series were averaged within each macro-region, yielding one EVC and one VVC PSI time series per participant and condition. PSI time series were smoothed with a 30-ms FWHM Gaussian kernel and analyzed from −0.2 to 1.0 s relative to target onset. The −0.2 s lower bound was chosen to avoid neural responses associated with cue offset. Predictable-and uninformative cue PSI time series were compared using paired-sample cluster-based permutation tests across time, with a cluster-forming threshold of p < 0.01, 5,000 permutations, and a cluster-level alpha of 0.05.

To determine whether target-specific representations emerged earlier after predictive cues, we estimated the onset of significant above-zero PSI separately for predictive and uninformative cue conditions. For each condition, onset was defined as the first time point of the earliest cluster in which PSI was significantly greater than zero. Statistical significance was assessed with one-sided cluster-based permutation tests. We then compared onset latencies using a one-sided bootstrap test. Participants were resampled with replacement 1,000 times, onset latencies were re-estimated for each resample using 2,000 permutations, and the onset difference was computed as uninformative minus predictive cue onset. A positive onset difference indicated earlier emergence of target-specific representations in the predictable-cue condition. The bootstrap distribution was summarized using the median and 95% interval, and the one-sided p-value was calculated as the proportion of bootstrap samples with onset differences less than or equal to zero.

We also performed a cross-phase PSI analysis to test whether pre-target activity reinstated target-specific patterns expressed after target onset. Using the same trial-to-template procedure described above, spatial activity patterns during the pre-target period were correlated with post-target target-specific templates. Cross-phase PSI was defined as the difference between the similarity to the matched-target template and the similarity to the alternative-target template, such that positive values indicated that pre-target activity resembled the subsequently expressed target-specific representation. PSI was averaged across post-target time intervals, showing significant differences between predictable-and uninformative-cue conditions in the time-resolved PSI analysis, yielding a pre-target PSI time series from −0.40 to −0.10 s. Statistical significance was assessed using cluster-based permutation tests, testing PSI against zero separately for each condition and directly comparing the predictable- and uninformative-cue conditions.

### Neural population trajectory analysis

To characterize how prediction altered the temporal evolution of population-level neural states, we performed a neural state-space trajectory analysis, following established population-dynamics approaches in which the activity pattern across a neural population at each time point defines a point in a low-dimensional state space, and its evolution over time defines a neural trajectory^33–35,51^. For each ROI, vertex-wise source activity was extracted from predictive- and uninformative-cue trials with the same target category and baseline-corrected relative to the pre-cue interval (−2.30 to −2.02 s).

To prevent differences in trajectory geometry from being driven by unequal trial counts, predictive and uninformative trials were matched within each run × cue-modality combination. Within each matched cell, an equal number of trials was randomly sampled from the two conditions, and this balancing procedure was repeated 50 times. For each repeat, condition-specific mean population responses across all matched cells and time points from −1 to 1 s relative to target onset were jointly submitted to PCA, such that predictive and uninformative trajectories were represented within a common neural state space. Vertex activity was standardized before PCA. Trajectory metrics were quantified in the first five PCs, consistent with previous MEG state-space analyses using a five-dimensional PCA representation of dominant population dynamics^36,52,53^.

We focused on trajectory dynamics during the pre-target interval from −0.40 to −0.10 s. Trajectories were smoothed with a 30-ms FWHM Gaussian kernel before computing movement metrics. Total path length was defined as the cumulative Euclidean distance between consecutive population states over the analysis interval, indexing the overall extent of state-space evolution^36,54^. This definition builds on previous neural state-space approaches that quantify moment-to-moment trajectory movement using the Euclidean distance between adjacent states in PCA space^36^. Step-distance variability was defined as the standard deviation of these consecutive Euclidean distances, indexing the consistency of moment-to-moment changes in population state. Both measures were calculated separately within each matched cell and balancing repeat before being averaged across cells and repeats. Predictive and uninformative conditions were compared using paired sign-flip permutation tests with 5,000 permutations.

For trialwise neural–behavioral analyses, trajectory measures were estimated using leave-one-run-out cross-fitting. For each held-out run, the PCA space was independently defined from the remaining runs using target-matched predictive and uninformative trials, balanced within each run × cue-modality combination. Each trial in the held-out run was projected into this independently defined state space, and single-trial trajectory metrics were computed and averaged across 50 balancing iterations.

### Trial-resolved neural–behavioral and cross-neural coupling

To examine whether trial-to-trial fluctuations in neural activity were related to response time, we fitted linear mixed-effects models predicting log-transformed response time from trial-wise neural measures. Consistent with the preceding neural analyses, these analyses were restricted to trials with target categories that were predictable in the predictive-cue condition and included trials from both the predictive-cue and uninformative-cue conditions. Neural predictors were standardized within participants before model fitting.

Separate models were fitted for each neural predictor, with cue condition included as a covariate and participant included as a random intercept:

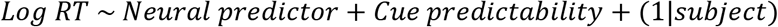

An additional model included motor beta lateralization and visual PSI simultaneously to estimate their independent associations with response time while controlling for cue condition.

Finally, multilevel mediation analyses were used to test whether relationships between neural activity and behavior were statistically consistent with indirect effects operating through other neural signals. Analyses were implemented in R using the multilevelmediation package^55^, with indirect effects evaluated using participant-level bootstrap resampling.

### DDM modeling and analyses

We used the drift-diffusion model (DDM) framework to test whether trial-by-trial neural measures associated with behavioral facilitation were related to latent decision parameters that jointly determine response speed. A hierarchical model-fitting approach was implemented using the HDDM package^56^. For each neural measure, we examined its associations with drift rate, decision threshold, and non-decision time, testing whether trial-by-trial variation in sensory, motor, or population-dynamics measures was associated with distinct components of the decision process. Analyses were restricted to uninformative-cue trials, in which living and non-living targets occurred equally often. Predictable-cue trials were excluded because they contained only one target category per participant, leaving insufficient trials to estimate one of the response boundaries. One participant was additionally excluded from the DDM analyses because a technical issue resulted in only 60 trials per block, leaving too few uninformative-cue trials for reliable model estimation. Non-living responses were coded as response = 1, and living responses were coded as response = 0. Neural predictors were z-scored within participant before model fitting.

Prior to neurocomputational modeling, we fitted the behavioral data to a baseline drift-diffusion model. The model included four parameters: drift rate, boundary separation, non-decision time, and starting-point bias (*v*, *a*, *t*, and *z*), which were estimated at both the group-level and individual levels using hierarchical methods.

Following our hypothesis-driven analysis plan, each neural predictor was tested against the three decision parameters (*v*, *a*, and *t*) that could plausibly account for faster responses. For category-specific neural measures, predictors entered into drift-rate models were coded directionally, such that stronger target-consistent neural evidence reflected greater accumulation toward the corresponding response boundary. Neural measures without an intrinsic response direction, including population-trajectory metrics, were entered as continuous unsigned predictors. For boundary-separation and non-decision-time models, all neural predictors were entered as unsigned measures. For both baseline and neural models, sampling was performed for 5,000 iterations, with 2,000 burn-in samples, and a thinning factor of 5. Model convergence was assessed by fitting five independent chains and computing Gelman-Rubin statistics for model parameters (all model parameters *R*^^^< 1.05).

## Data and materials availability

The behavioral and MEG data and analysis code supporting the findings of this study will be made publicly available on Zenodo upon publication.

## Author contributions

L.Z. and C.A. designed research; L.Z. and M.Y. performed research; L.Z. analyzed data; L.Z. wrote the first draft of the paper; and L.Z., M.Y., H.Z., M.M., and C.A. reviewed and edited the paper.

---

**Figure S1.**
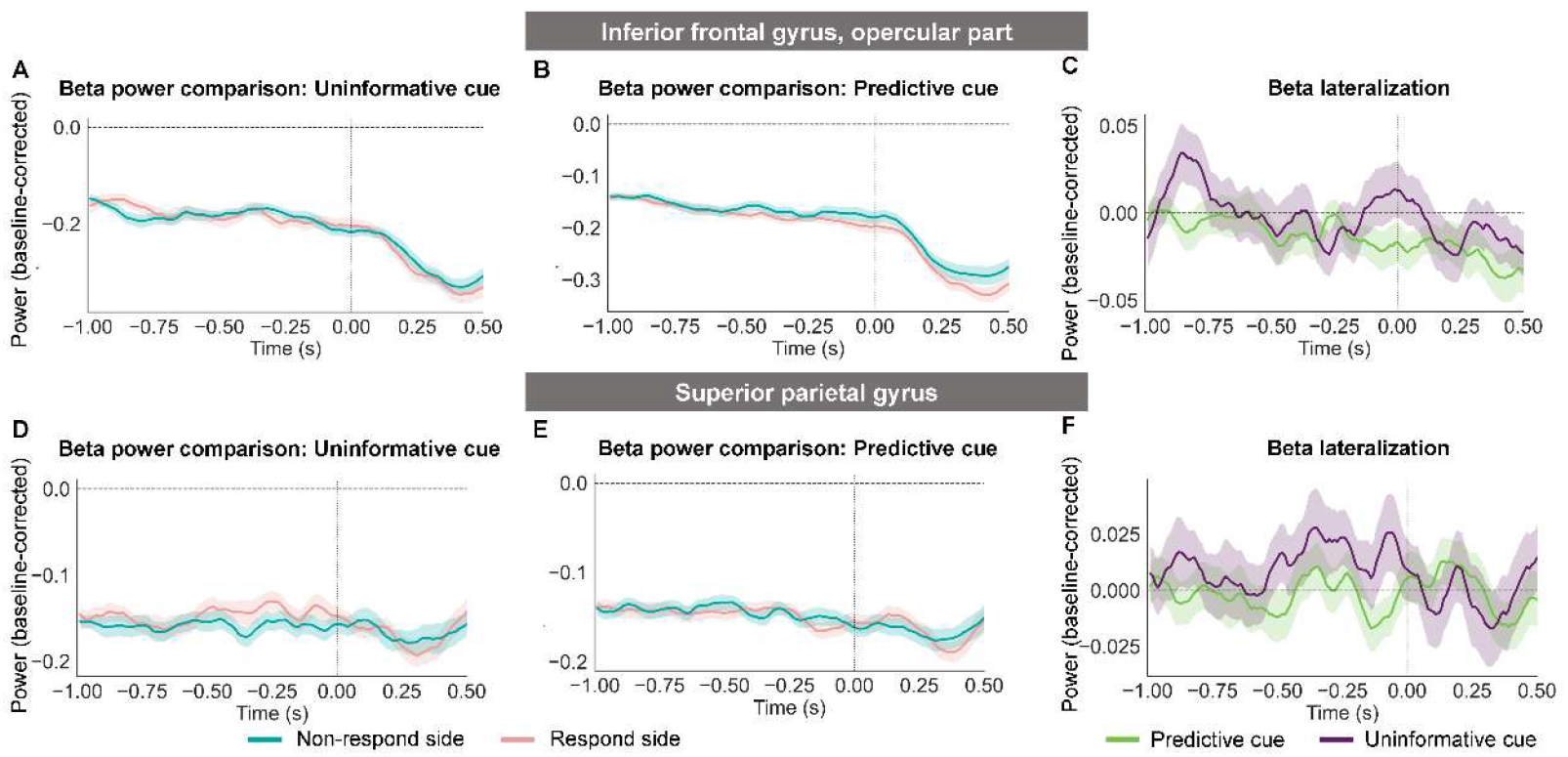
Control analyses of beta suppression in inferior frontal and superior parietal regions. A–C, Beta power in the inferior frontal gyrus, opercular part, following uninformative (**A**) and predictive (**B**) cues, and the corresponding beta lateralization time courses (**C**). **D–F,** Corresponding analyses in the superior parietal gyrus following uninformative (**D**) and predictive (**E**) cues, with beta lateralization shown in **F**. No reliable cue-condition differences in beta lateralization were observed in either control region. Shaded envelopes indicate SEM. Time 0 denotes target onset. SEM, standard error of the mean.

**Figure S2.**
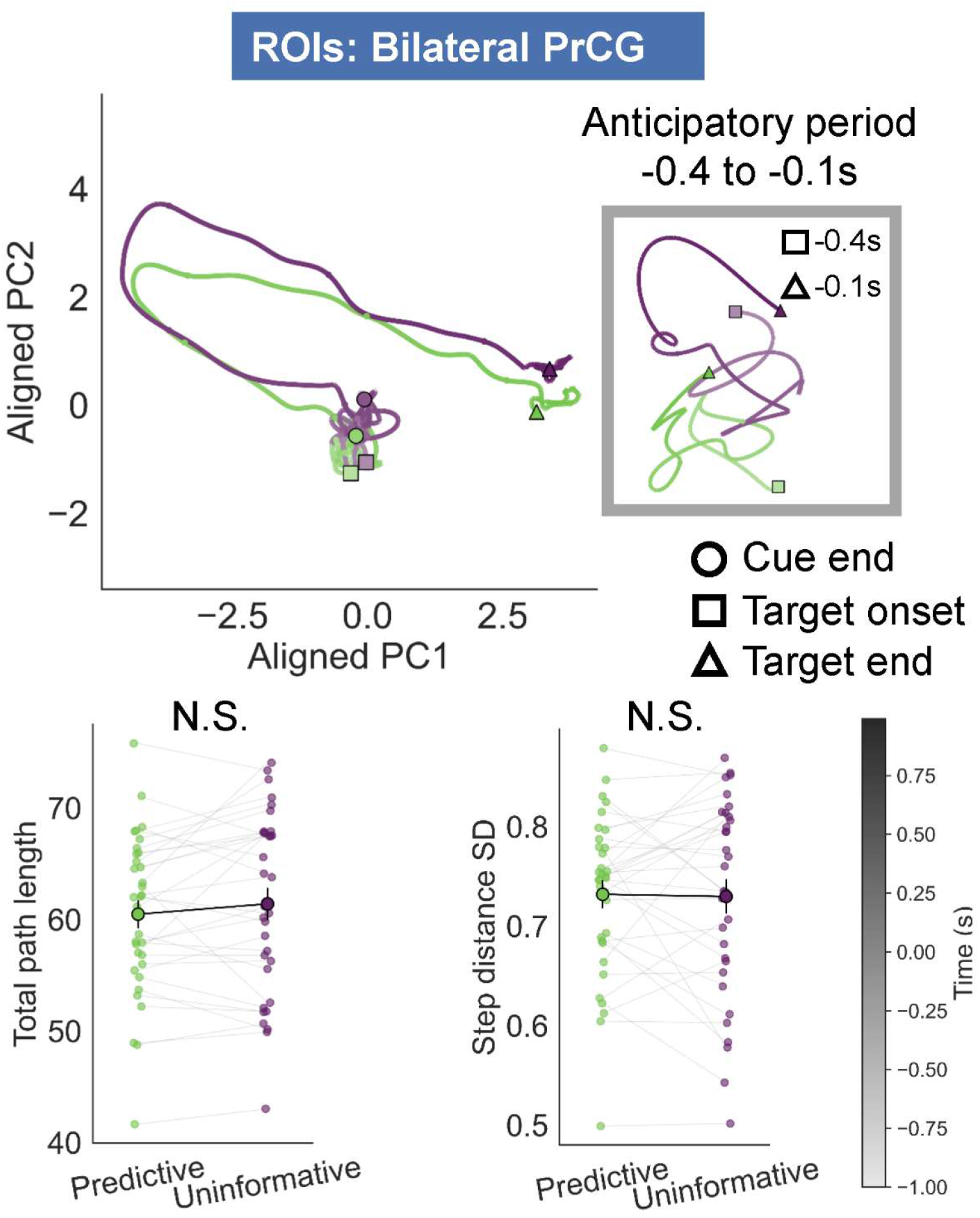
Predictive cues did not reliably alter anticipatory population dynamics in motor cortex. Population trajectories through a low-dimensional neural state space in bilateral PrCG following predictive and uninformative cues. The inset shows trajectories during the anticipatory period from −0.40 to −0.10 s. Symbols mark cue offset, target onset, and target offset. Neither total path length nor trajectory variability differed reliably between cue conditions. PrCG, precentral gyrus; PC, principal component; N.S., not significant.

